# Novel Roles of G-quadruplexes on Enhancers in human chromatin

**DOI:** 10.1101/2021.07.12.451993

**Authors:** Yue Hou, Yan Guo, Shanshan Dong, Tielin Yang

## Abstract

G-quadruplexes (G4), stable four-stranded non-canonical DNA structures, are highly related to function of promoters and initiation of gene transcription. We found that G4 structures were also enriched in the enhancers across different cell lines. However, the relationship between G4 structures and enhancer activity remains unknown. Here, we proved that G4 structures on enhancers lead to the re-positioning of nucleosomes create nucleosome depleted regions (NDRs). Moreover, stable NDRs and special secondary structures of G4 help enhancers to recruit abundant TFs to co-bind, especially for architectural proteins including CTCF, RAD21, and SMC3. These architectural proteins, which play critical roles in the formation of higher-order chromatin organization, further influenced the chromatin interactions of G4 enhancers. Additionally, we revealed that G4 enhancers harbored significantly higher enrichment of eQTLs than typical enhancers, suggesting G4 enhancers displayed more enhancer regulatory activity. We found that most super enhancers (SEs) contain G4 structures. Even though the enrichment of chromatin accessibility and histone modifications around G4-containing SEs are not significantly higher than those around other SEs, G4-containing SEs still possess much more TFs across different cell lines. According to these results, we proposed a model in which the formation of G4 structures on enhancer exclude nucleosome occupancy and recruit abundant TFs which lead to the stable chromatin interaction between G4 enhancers and their target genes. Because of the relevance between G4 structures and enhancers, we hypothesized that G4 structures may be a potential markers indicating enhancer regulatory activity.

## Introduction

Although DNA double-helix structure is the predominant configuration inside the cells, a wide variety of non-canonical secondary structures, including G-quadruplex (G4), R-loop, H-DNA, Z-DNA et al., have been detected both *in vitro* and *in vivo* (Georgakopoulos-Soares et al., 2018). G4 structure is a stable secondary structure formed by square planes where four guanines located in the same plane stabilized by a monovalent cation (Varshney et al., 2020). Taking advantage of polymerase stalling at G4s, G4-seq identified more than 700,000 DNA G4 structures *in vitro* (Chambers et al., 2015). To detect DNA G4s in the chromatin of human cells, G4 antibody called BG4 was used in chromatin immunoprecipitation followed by high-throughput sequencing (G4 ChIP-seq) which confirmed more than 10,000 high confidence G4 structures in specific cell lines (Hansel-Hertsch et al., 2016; Hansel-Hertsch et al., 2018). Most recently, an artificial 6.7 kDa G4 probe (G4P) protein, which binds G4s with high affinity and specificity, was applied to capture G4s in living human with the ChIP-seq technique and detected up to >123,000 G4P peaks (Zheng et al., 2020).

The early work about G4s mainly focused on their roles in telomeres and telomerase (Henderson et al., 1987; Zaug et al., 2005). A number of subsequent studies further demonstrated that G4 structures substantially participate in the regulation of gene expression in either direction (Du et al., 2009; Renciuk et al., 2017). Initially G4 motifs were found to be highly enriched in the gene promoter regions by computational predictions (Huppert and Balasubramanian, 2007). Several followed studies extended this result to a chromatin context by revealing the enrichment of G4s in promoters *in vivo* (Hansel-Hertsch et al., 2016; Zheng et al., 2020). In the first exon of *hTERT*, the formation of G4 structure led to the high expression level of *hTERT* (Li et al., 2017). Studies inside gene bodies showed a substantial inhibitory effect of G4 structures in the elongation of RNA polymerase, resulting in low expressed genes (Agarwal et al., 2014; Holder and Hartig, 2014). Benefiting from the special secondary structures, G4s have high affinity for different transcription factors (TFs) and specific proteins (Hansel-Hertsch et al., 2020; Li et al., 2020; Mao et al., 2018). Because of this character, G4 structures can regulate DNA looping and gene expression by attracting Yin and Yang 1 (YY1) (Li et al., 2020). Overall, the weight of evidence from previous studies showed the biological function of G4s in the gene-proximal regions. Nevertheless, the relevance of G4s and promoter-distal regulatory elements, such as enhancers, remains unclear.

Enhancers are genomic regulatory elements that regulate gene expression regardless of location and orientation through bound by abundant TFs. It is important for enhancers to recruit various TFs which further lead to transcription of enhancers (Kim et al., 2010; Sartorelli and Lauberth, 2020). It is known that RNA polymerase II (RNA Pol II) on enhancers transcribes bi-directionally and generates non-coding RNAs (eRNAs) (De Santa et al., 2010; Kim et al., 2010). Consequently, eRNAs are highly related to the activity of enhancers (Hah et al., 2013; Li et al., 2013). Intriguingly, the bi-directionally transcription property was observed in both enhancers and promoters (Core et al., 2014; Hah et al., 2013; Melgar et al., 2011). It is inferred that enhancers and promoters independently recruit TFs, but have to cooperate with each other to achieve efficient transcription (Li et al., 2016). Through analyzing TCGA RNA-seq data, Chen and Liang suggested that well-positioned nucleosomes in super enhancers were involved in the regulation of enhancer transcription (Chen and Liang, 2020). However, the determinants of nucleosome positioning and TF binding at active enhancers remains elusive.

Due to the close relationship between G4 structures and TF binding, we hypothesized that the formation of G4 structures at enhancers can also regulate the activity of enhancers by affecting nucleosome repositioning and TF binding. In this study we methodically explored the relationship between G4s and enhancers. G4P and BG4 ChIP-seq data generated by previous studies were used to call G4 peaks that were regarded as G4 structures (Mao et al., 2018; Zheng et al., 2020). We revealed that G4 structures are highly related to the chromatin structure and TF binding on enhancers which further influence enhancer transcriptional events. Using the recently developed powerful ChIA-pet technology, we investigated that how G4s act their regulation function in the higher-order chromatin context. Finally, we propose that G4 structures act as markers of enhancer regulatory activity. These findings expand our knowledge on the function of G4 structures.

## Results

### G-quadruplexes exclude nucleosome occupancy on enhancers

A number of studies have revealed the roles of G4 structures on promoters (Renciuk et al., 2017; Shen et al., 2021; Varshney et al., 2020; Zheng et al., 2020). However, there are many G4 structures located in the gene-distal regions and highly enriched in enhancers regions across different cell lines including A549, HEK293T, HeLa-S3 and K562 cell lines (Figure S1A-B). Because of high abundance of GC-rich regions, promoters always contain stable G4 structures. Enhancers are much less GC-rich than promoters, nonetheless, they were still rich in G4 structures (Figure S1). To find out whether these G4 structures formed in enhancer regions were functional, we divided the enhancers into different groups: G4-associated enhancers and typical enhancers. In line with previous research, H3K4me1 ChIP-seq peaks overlapped with DNase-seq peaks were designated as enhancers (Doni Jayavelu et al., 2020). We termed the enhancers overlapped with G4 peaks as G4-associated enhancers (G4 enhancers, Figure 1A–B); the rest of enhancers were termed as typical enhancers. G4P and BG4 ChIP-seq data generated from previous studies were used to achieve the accurate positions of G4 peaks *in vivo* (Mao et al., 2018; Zheng et al., 2020).

**Figure 1.**
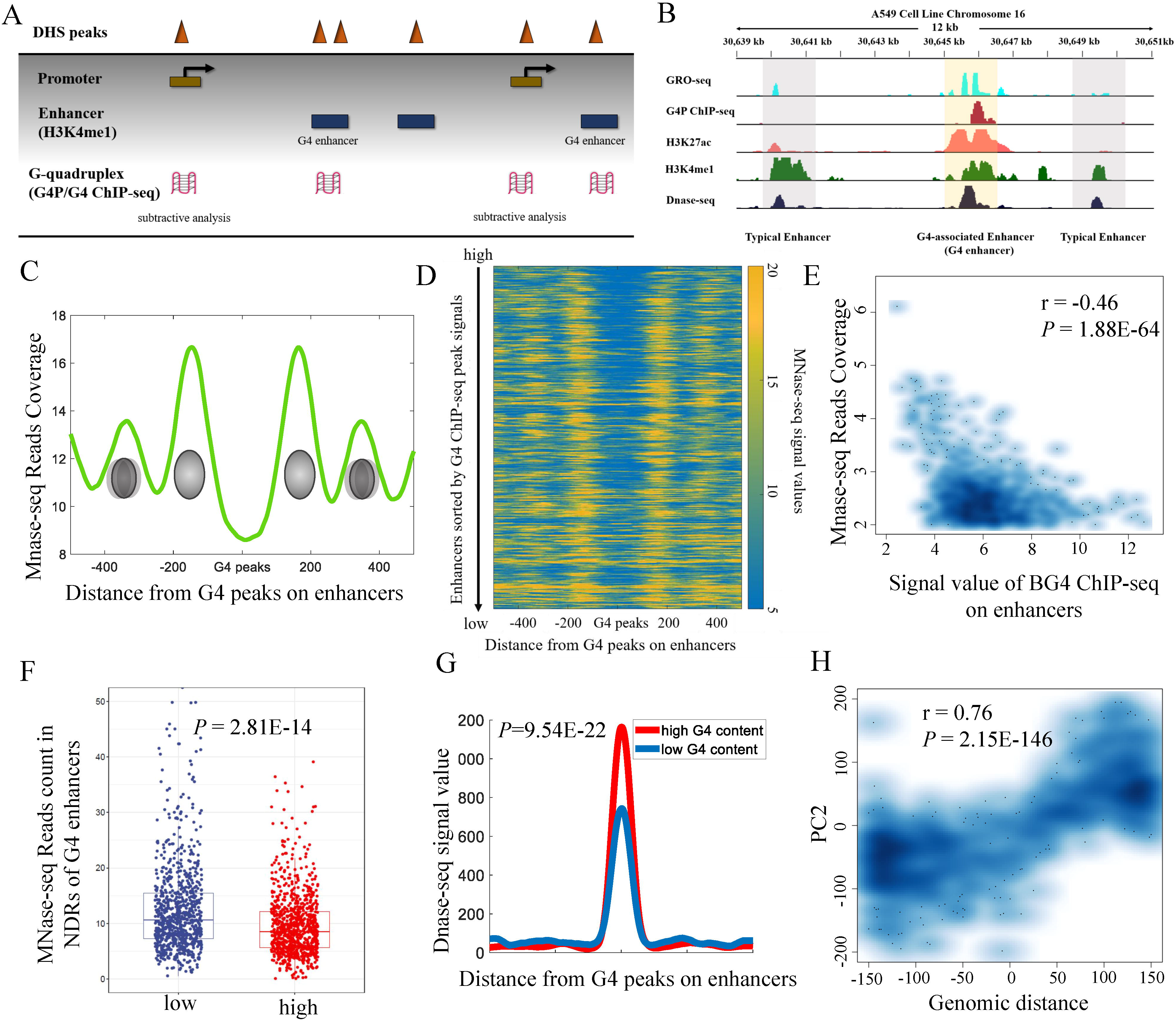
G4 structures maintain stable NDRs on enhancers. (A) Definition of G4 enhancers. Gene distal H3K4me1 ChIP-seq peaks overlapped with DHS peaks were defined as enhancers. G4 enhancers were designated as enhancers overlapped with G4 peaks. (B) An IGV screen shot of G4 enhancers in the A549 cell line. (C) Occupancy of nucleosome around G4 structures of enhancers in the K562 cell line. (D) Heatmap of nucleosome signals around G4 structures of enhancers in the K562 cell line. All rows were sorted according to the G4 peak signal values. (E) G4 structures are negative correlated with MNase-seq read counts on enhancers. (F) High G4 structures lead to deeper NDRs on enhancers (Student's t-test *P*-value = 2.81E-14). (G) high G4 enhancers displayed higher chromatin accessibility than low G4 enhancers. (H) Principe-component analysis (PCA) of nucleosome positioning on the G4 peaks. The correlation of PC2 with the genomic distances between first GG-run and the middle site of G4 peaks.

It is notable that the well-positioned nucleosomes in enhancers directly regulate enhancer activity (Andersson and Sandelin, 2020; Chen and Liang, 2020). Herein, we found it is G4 structures that shape the nucleosome pattern on enhancers in the K562 cell line (Figure 1C–F). Through calculating MNase-seq reads coverage on enhancers, we found that nucleosomes are excluded from G4 peaks, and thus formed nucleosome depleted regions (NDR, Figure 1C–D). Additionally, nucleosomal arrays were found in the upstream and downstream regions of G4 peaks (Figure 1C–D), indicating that the formation of G4 structures lead to the re-position of nucleosomes in enhancer regions. Furthermore, G4 signal values on enhancers were negatively correlated with the MNase-seq reads coverage (Figure 1D–E, Pearson's R = −0.46, *p* < 1.88e-64), suggesting that in result in high nucleosome occupancy and fuzzy NDRs on enhancers (Figure 1D–E). We further divided G4 enhancers into two groups: high G4 enhancers and low G4 enhancers according to the G4 peak values on the enhancers. G4 enhancers were averaged to two groups, those with high G4 signals called high G4 enhancers and those with low G4 signals called low G4 enhancers. The low G4 enhancer group harbored significantly more MNase-seq read counts than the high G4 value group (Student's t-test *P*-value = 2.81E-14, Figure 1F). Due to the depleted nucleosome occupancy, the high G4 enhancers have more accessible DNA than low G4 enhancers (Student's t-test *P*-value = 7.57E-94, Figure 1G). This prominent association between G4 peaks and NDRs suggested that unstable G4 structures result in high nucleosome occupancy and fuzzy NDRs.

To characterize the major patterns of nucleosome binding around G4 structures on enhancers, we performed principal-component analysis (PCA) on the signals (Figure S2A). The first two principal component (PC) explained more than 82% of the total variations. PC1 reflected the nucleosome signal values around G4 structures on enhancers (Figure S2B, Pearson's R = 0.99, *p* < 2.53e-266). PC2 was independent of nucleosome signal values (Figure S2C-D). PC2 represented the phase of nucleosome around G4 structures on enhancers (Figure S2E-F). Interestingly, the genomic distance between the first GG-run and the center of G4 peaks displayed positive correlated with PC2 (Figure S2D, Pearson's R = 0.76, *p* < 2.15e-146), suggesting that G4 structures not only determine the intensity of nucleosome, but also determine the phase of nucleosome on enhancers.

### G-quadruplexes amplify enhancer regulatory activity by influencing TF binding

NDRs in regulatory elements ensure that DNA is accessible to the TFs. To check whether G4 structures were potentially influencing TF binding on enhancers, we further calculated the TF ChIP-seq peak counts around the G4 enhancers. In the four cell lines, G4 enhancers contain much more TF ChIP-seq peaks than typical enhancers (Figure 2A–D, Student's t-test *p* < 3.10e-128, 5.77e-165, 9.22e-112 and 6.98e-177 for A549, HEK293T, HeLa-S3, and K562 cell lines, respectively). To find out which transcription factors are specifically recruited by the G4 structures on enhancers, we performed Fisher's exact test on each TF ChIP-seq in the enhancer regions and used the Benjamini-Hochberg method for correction. We selected the TFs, for which fold enrichment > 5 and q-value <1e-10, as G4 enhancer-enriched TFs. The results indicated that most G4 enhancer-enriched TFs is cell type specific, even though some G4 enhancer-enriched TFs are conserved across different cell lines (Table S1-S4). A total of 16, 11, 52, and 98 TFs are significantly enriched on G4 enhancers in A549, HEK293T, HeLa-S3, and K562 cell lines, respectively (Table S1-S4). In A549 cell line, G4 enhancers harbored enriched CREB1 and CRTC2 binding sites (Table S1, fold enrichment = 62.20 and 142.14, respectively; log Q-value =− 283.05 and −222.38, respectively). It has been reported that the CREB coactivator CRTC2 promotes tumor growth, and can be one potential therapeutic target of non-small cell lung cancer (Rodon et al., 2019). Herein, our results suggested in the non-small cell lung cancer cell line (A549 cell line) it is G4 structures that recruit CREB1 and CRTC2 to bind on enhances.

**Figure 2.**
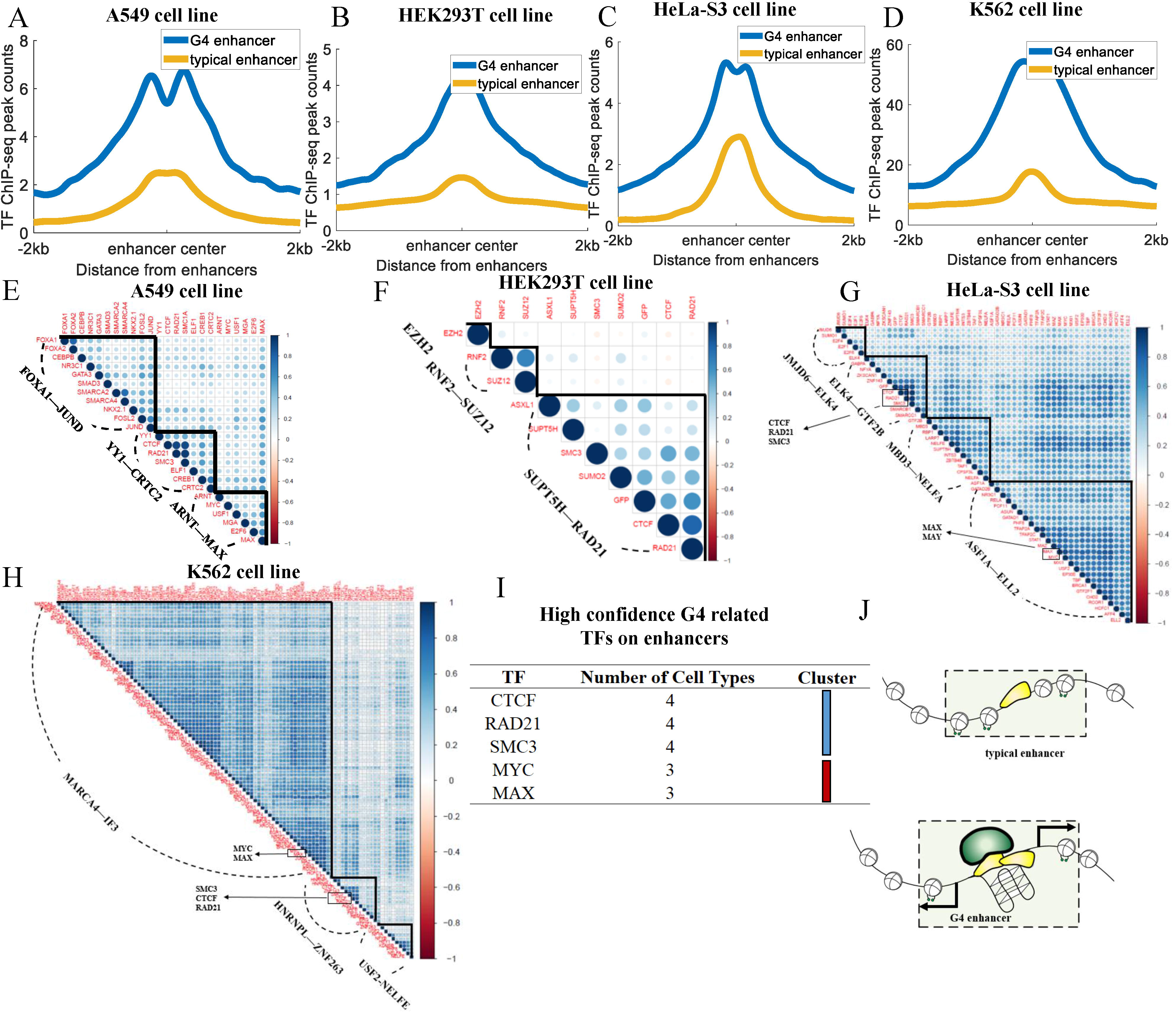
G4 structures are highly related to the binding of TFs around the enhancers. (A-D) Comparison of the distribution of transcription factor binding in enhancer region in different cell lines; (E-H) Heat map of transcription factor co-binding recruited by G4s on enhancer region in different cell lines; (I) Conserved high confidence G4 related TF clusters on enhancers; (J) Stable formation of G4s in enhancers exclude the nucleosomes to form NDRs. Moreover, the special secondary structure of G4s was beneficial for enhancers to recruit different TFs.

We used hierarchical cluster analysis of the enriched TFs on G4 enhancers and found that G4s can attract different clusters of TFs to co-bind on the enhancer regions (Figure 2E–H). Two types of clusters are conserved in different cell lines (Figure 2I). One cluster includes three architectural proteins including CTCF, SMC3, and RAD21; and the other cluster includes MYC and MAX (Figure 2I). It has been reported by previous research that CTCF and cohesin proteins directly induce the enhancer– promoter interactions (Li et al., 2015; Rao et al., 2017; Tang et al., 2015). Herein, we found that the formation of G4s help enhancers to recruit these architectural proteins. Both MYC and MAX are significantly enriched in G4 enhancers, and they can form heterodimer protein to activate transcription (Cascon and Robledo, 2012).

It has been proved that enhancer transcription is a key indicator of enhancer regulatory activity (Kim et al., 2010; Sartorelli and Lauberth, 2020). We speculated that stable NDRs and high abundant TF bindings in G4 enhancers directly facilitate the transcription events on enhancers. To check the hypothesis, we calculated GRO-seq signals on different types of enhancers. We adopted the global nuclear run-on sequencing (GRO-seq) data, which are capable of capturing 5′-capped RNAs from active transcriptional regulatory elements with high accuracy (Danko et al., 2015), to determine the transcriptional events on enhancers. We found that GRO-seq signals on G4 enhancers were significantly higher than those on typical enhancers in the four cell lines including A549, HEK293T, HeLa-S3, and K562 cell line (Figure S3A-D) suggesting that G4 structures on enhancers can promote the transcription of enhancer RNAs (eRNAs).

Taken together, our results support a model that the stable formation of G4s exclude the nucleosomes in enhancers to form NDRs which can influence the enhancer transcription by recruiting different TFs (Figure 2J).

### G-quadruplexes are related to the long-range chromatin interactions by recruiting chromatin architectural protein

According to previous studies, enhancer transcripts are involved in maintaining chromatin loops in different ways (Bouvy-Liivrand et al., 2017; Lam et al., 2014; Yang et al., 2016). Furthermore, the folding of G4 structures is highly associated with RNA polymerase II (Pol II) (Shen et al., 2021). Since G4 enhancers are rich in transcriptional signals, we compared the interaction reads from RNA Pol II ChIA-pet in the G4 enhancer groups with those in typical enhancer groups. It can be seen that in all four cell lines G4 enhancers contain much more RNA Pol II ChIA-pet reads (Figure 3A–D).

**Figure 3.**
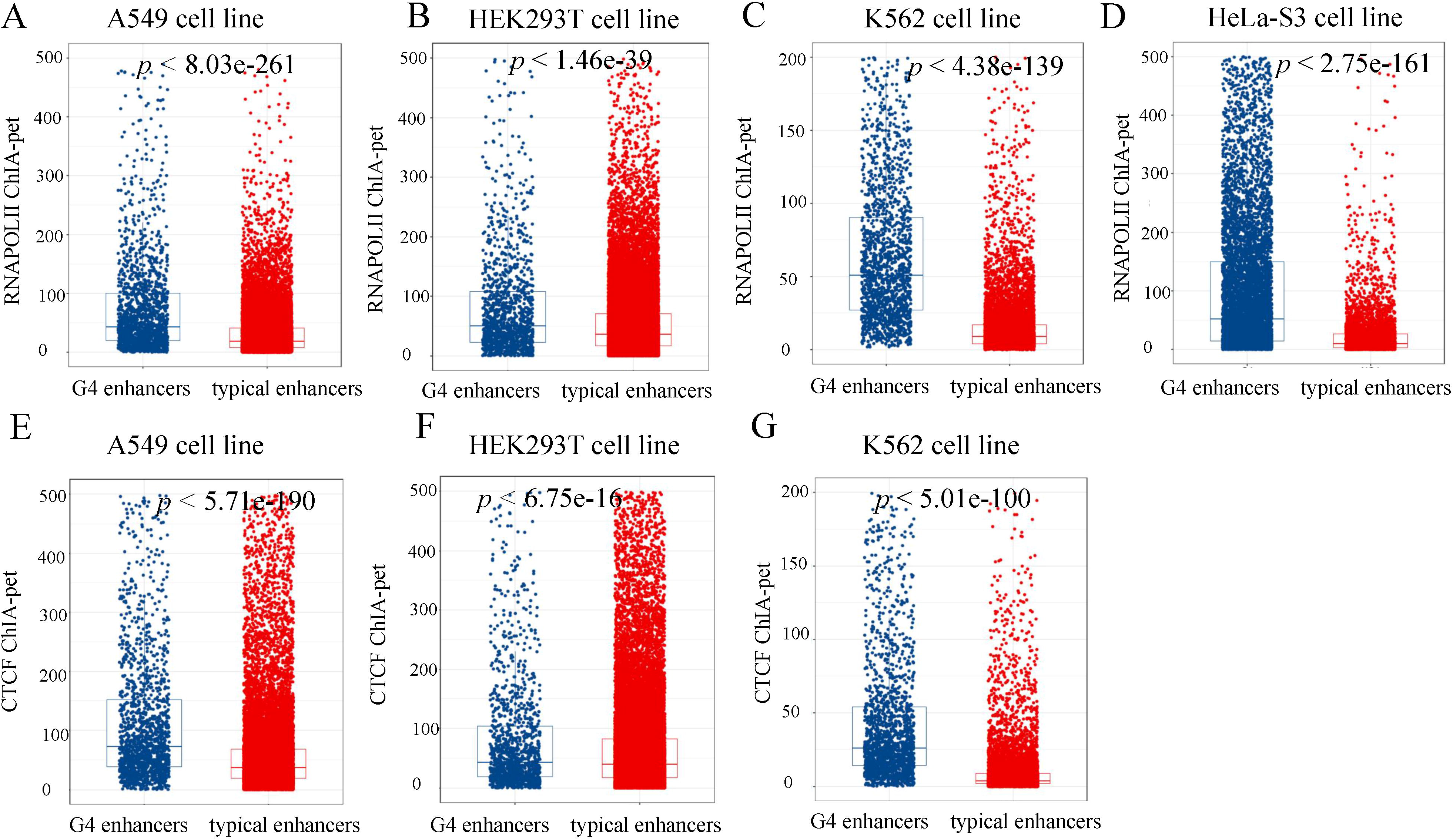
Comparison of ChIA-pet reads between G4 enhancers and typical enhancers. (A-D) G4 enhancers have more RNA POLII ChIA-pet reads than typical enhancers in the A549, HEK293T, K562 and HeLa-S3 cell lines, respectively. (E-F) G4 enhancers have more CTCF ChIA-pet reads than typical enhancers in the A549, HEK293T, and K562, respectively.

Architectural proteins including CTCF, SMC3, and RAD21, play key roles in mediating long-range chromatin interactions (Hong and Kim, 2017; Rao et al., 2017; Vietri Rudan et al., 2015). As CTCF, SMC3, and RAD21 are conserved TF clusters binding in G4 enhancer regions (Figure 2), we conclude that G4 structures may also influence the chromatin loops by recruiting these architectural proteins. In line with our hypothesis, G4 enhancers also possess more CTCF ChIA-pet reads (Figure 3E–G). Since ENCODE database only publish CTCF ChIA-pet of three cell lines including A549, HEK293T, and K562 cell lines, only CTCF ChIA-pet reads in these cell lines were shown. Taken together, the results suggest that G4 structures can activate enhancer transcription and recruit CTCF to mediate chromatin interactions.

### The folding of G4 structures on enhancers is associated with super enhancers in human chromatin

Core super enhancers (SEs) always generate stable eRNA and possess abundant TF binding (Chen and Liang, 2020). Because of the close relationship between G4 structures and enhancer transcription, we wondered whether the folding of G4 structures on enhancers is associated with SEs. We used the standard RANK ORDER of SUPER ENHANCERS (ROSE) algorithms to calculate the accurate position of SEs (Whyte et al., 2013). Strikingly, we found that most SEs were overlapped with G4 peaks (Figure 4A–D). Compared with other SEs, G4-containing SEs display no significant difference in H3K4me1 ChIP-seq signals and DNase-seq signals (Figure S4). However, G4-containing SEs harbor much more TF ChIP-seq peaks than other SEs (Figure 4E–H). Since there is no significant difference of active histone modification signals and DNA accessible signals between G4-containing SEs and other SEs, the abundant TF ChIP-seq peaks on G4-containing SEs should be resulted from the formation of G4 structures.

**Figure 4.**
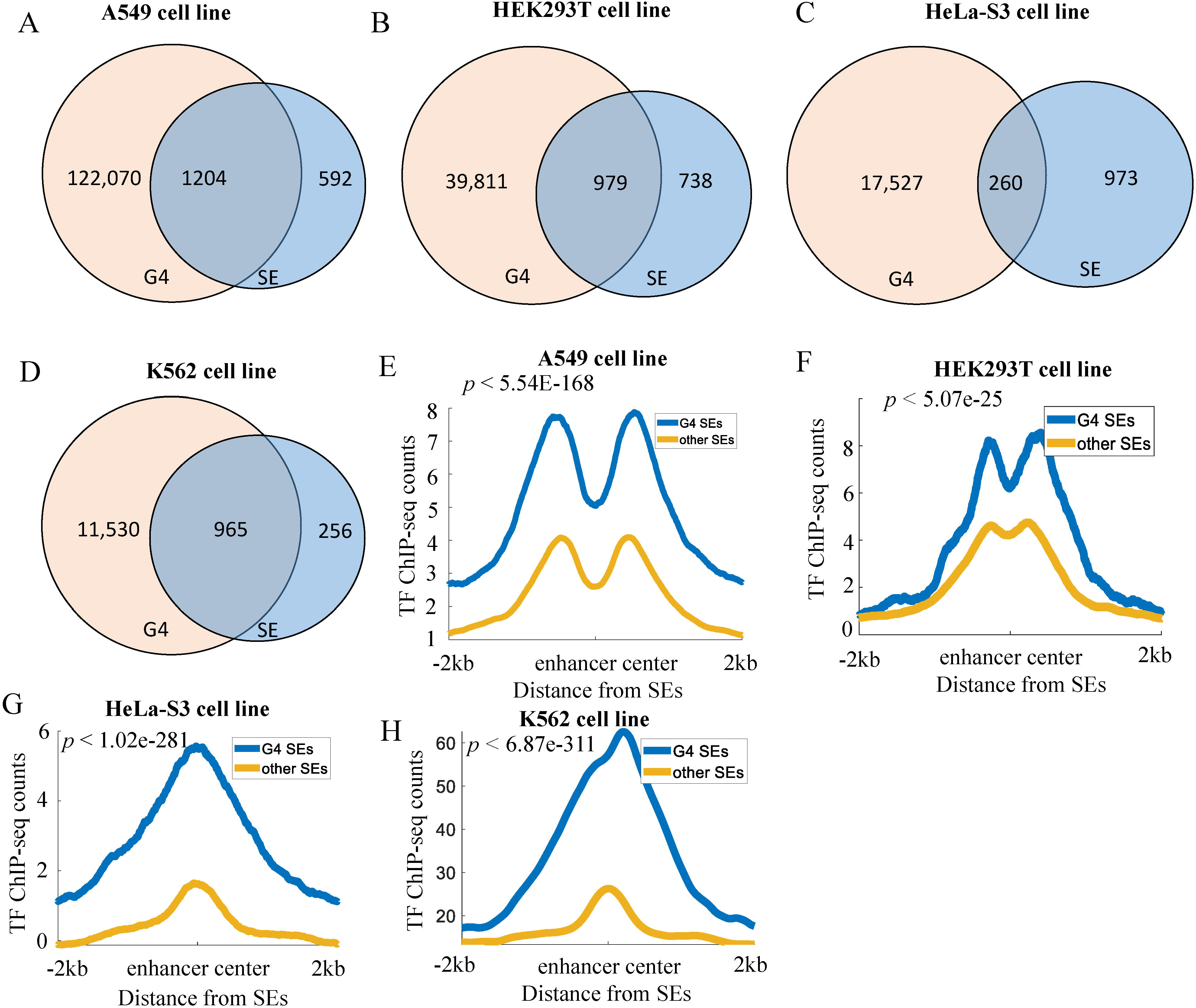
G4 peaks were highly enriched in the super enhancer regions. (A-D) Venn diagram of SEs and G4 peaks in the four cell lines. (E-H) Comparison of the TF ChIP-seq peak counts in G4-containing SEs and other SEs in the four cell lines.

### Disease risk variants are highly enriched in G-quadruplexes-associated enhancers

Genetic variations within regulatory genomic elements often associate with variation in expression of the linked target genes. As such, expression quantitative trait loci (eQTL) enrichment analysis can serve as an objective and quantitative metric to evaluate regulatory potential. We compared the frequencies of SNPs and eQTLs within G4 enhancers and typical enhancers to evaluate the regulatory potential of G4 structures (Figure 5). SNPs were selected from GWAS catalog database (*p* < 5e-8). And eQTL of lung, kidney, whole blood, and uterus were downloaded from GTEx eQTL database. To measure the enrichment of SNP and eQTL, we calculate the fold enrichment score of SNP and eQTL within the different types of enhancers relative to genome background.

**Figure 5.**
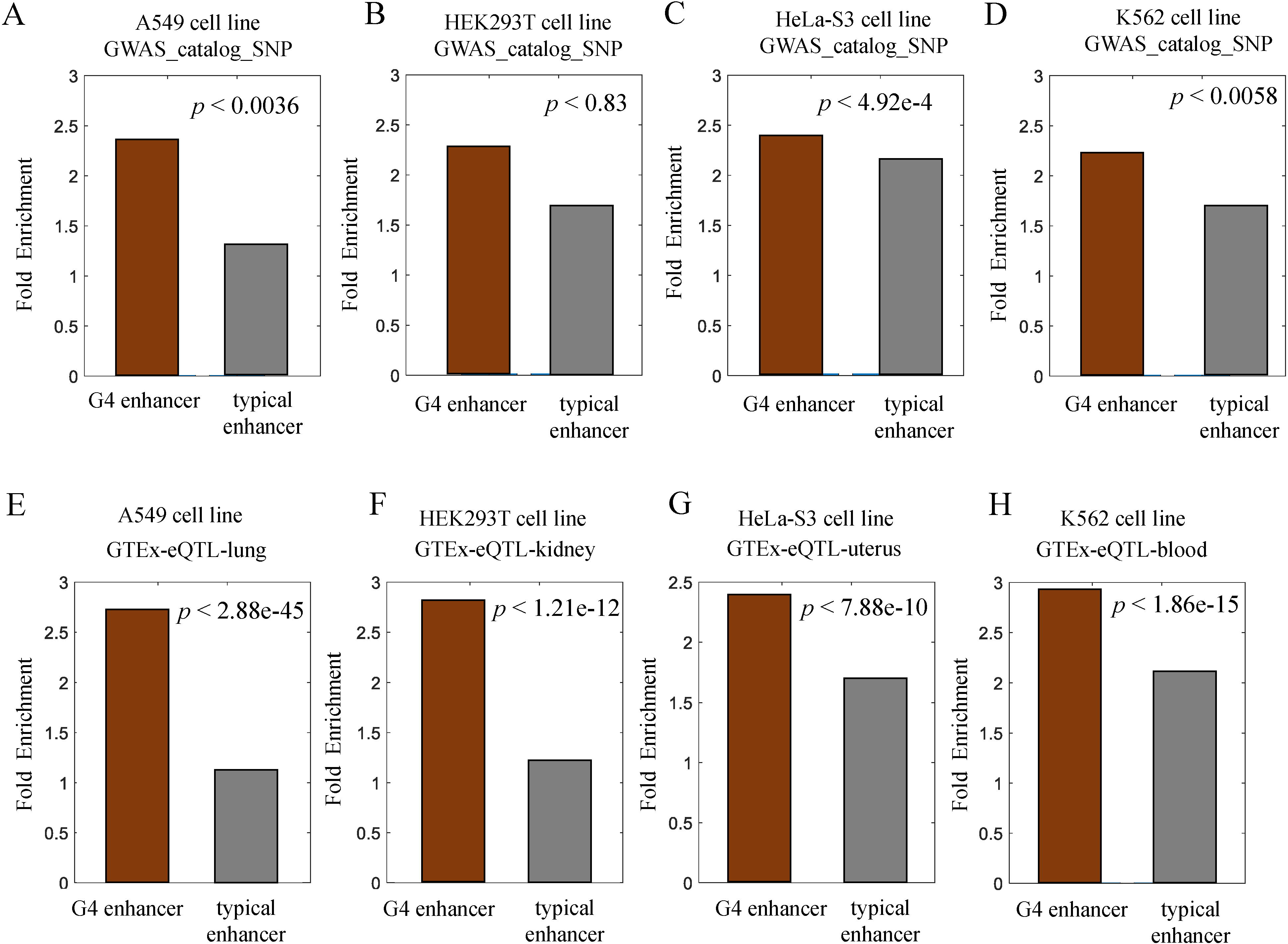
Enrichment of genetic variants associated with diseases and cell-type-specific gene expression in G4 enhancers in different cell lines. (A–D) Bar plot of GWAS SNP fold enrichment within G4 enhancers and typical enhancers in different cell lines. *P* values were calculated using Fisher's exact test. (E-F) Bar plot of eQTL fold enrichment within G4 enhancers and typical enhancers in different cell lines. *P* values were calculated using Fisher's exact test.

The GWAS SNP fold enrichment values for G4 enhancers in A549, HeLa-S3 and K562 are moderate higher than those in typical enhancers (Figure 5A–B, and D, Fisher's exact test, *p* < 3.6e−3, 4.92e-4, and 5.8e-3 for A549, HeLa-S3 and K562, respectively). Nevertheless, in HEK293T cell lines there is no significant difference of GWAS SNP fold enrichment between G4 enhancers and typical enhancers (Figure 5C, Fisher's exact test, *p* < 0.83). We found that in all four cell lines the fold enrichment values of eQTL for G4 enhancers are significantly higher than those for typical enhancers (Figure 5E–H, Fisher's exact test, *p* < 2.88e-45, 1.21e-12, 7.88e-10, and 1.86e-15 in A549, HEK293T, HeLa-S3, and K562 cell lines, respectively).

Taken together, our studies demonstrate that G4 enhancers are more enriched with genetic variants associated with diseases than typical enhancers, suggesting they may play a more important role in developmental control and mediating disease risks.

### G4 structures as potential markers of enhancer regulatory activity in human chromatin

Due to the relevance between G4 structures and enhancers, we wondered if G4 structures could be new markers of enhancer regulatory activity in the human chromatin. It has been proved that GRO-seq data can indicate the active transcriptional regulatory elements (Danko et al., 2015). We observed most, if not all, G4 structures located in non-coding regions displayed a high accumulation of GRO-seq signals in the four cell lines, with the signals found to gradually decline towards the flanking regions of G4 peaks (Figure 6A–D). Moreover, a double-peak phenomenon of GRO-seq signals, which is similar to those on transcriptional enhancers, appeared around the G4 peaks (Figure 6A–D). Our results showed that almost all gene promoter distal G4 peaks displayed transcriptional activities according to the GRO-seq signals. To exclude the influence of enhancers and genes, only G4 structures (individual G4 structures, iG4s) that are not overlapped with enhancers, promoters, or known genes, were retained. GRO-seq signals were significantly enriched in iG4 than enhancers predicted by H3K4me1 (Figure 6E–H). Besides, the counts of TF binding on these iG4s were also significantly higher than those on enhancers (Figure 6I–L), suggesting these iG4s were TF binding hubs even though they are not predicted enhancers and promoters. Taken together, our results suggested G4 structures can be potential markers of enhancer regulatory activity in human chromatin.

**Figure 6.**
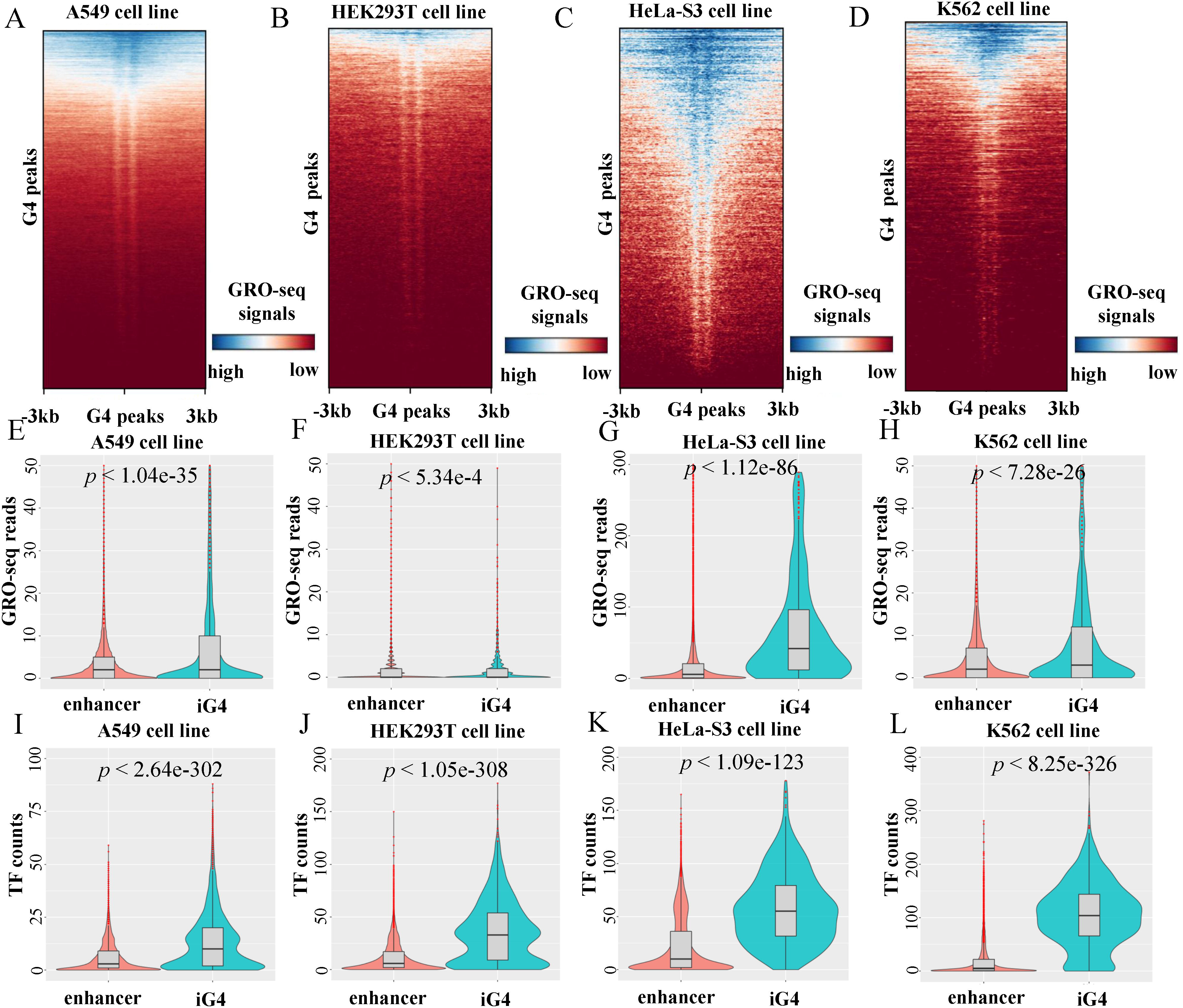
G4 structures as potential markers of enhancer regulatory activity. (A–D) Heatmaps of GRO-seq signals on promoter-distal G4 peaks in the four cell lines. (E-H) Violin plot of GRO-seq signals within enhancers and iG4s in different cell lines. *P* values were calculated using a two-sided Wilcoxon rank sum test. (I-L) Violin plot of TF binding counts on enhancers and iG4s in different cell lines.

## Discussion

There is a broad consensus that active enhancers recruit many TFs and generate non-coding transcripts which indicate the activity of enhancers (Li et al., 2016). Nevertheless, how TFs were recruited to the enhancer regions remains unknown. In this study, we characterized G4 structures on enhancers in the different human cell lines. Using both DNase-seq and histone modification data, we identified active enhancers in the four cell lines including the A549, HEK293T, HeLa-S3, and K562 cell lines (Figure 1). The dynamic, well-positioned nucleosomes enhancers directly regulate the activity of enhancers (Andersson and Sandelin, 2020; Chen and Liang, 2020). In this study, we proved that G4 structures on enhancers re-positioned nucleosomes leading to the NDR and well-positioned nucleosome arrays around G4 peaks (Figure 1). The results revealed that the formation of G4 structures on enhancers can potentially regulate enhancer activity through maintenance of the NDRs.

Because NDRs provide accessible DNA for specific TFs to bind on, we speculated that G4 structures on enhancers can attract certain TFs. Through enrichment analysis, we found G4 enhancers harbored different TF clusters in different cell lines (Figure 2). Among these TF clusters, CTCF, SMC3, and RAD21 were conserved in all the cell lines. Previous studies showed that SMC complex can extrude DNA at a high speed to form higher-order genome(Ganji et al., 2018; Wang et al., 2017). In the chromatin extrusion model, CTCF can stop the sliding of the SMC complex to form chromatin loop structures (Sanborn et al., 2015). Most chromatin loop anchors contain typical enhancer-promoter interactions. Our analysis suggested that G4 structures on enhancers may participate in the loop extrusion by recruit the architectural proteins including CTCF, RAD21, and SMC3 (Figure3). It can be seen that in all four cell lines G4 enhancers contain much more RNA POLII ChIA-pet reads (Figure 3A–D). Additionally, G4 enhancers also possess more CTCF ChIA-pet reads (Figure 3E–G).

eQTL enrichment analysis can serve as an objective and quantitative metric to evaluate regulatory potential. In line with our hypothesis, we found that eQTL were much more significantly enriched in G4 enhancers than typical enhancers, even though SNP enrichment in these two types enhancers are almost same (Figure 5). It suggested that G4 enhancers tend to have more regulatory activity. Intriguingly, we found that most SEs, if not all, possess G4 peaks (Figure 4). We found that G4 SEs have more TF binding sites than other SEs and this is not resulting from the DNA accessibility and histone modifications. Nonetheless, it remains unclear that if G4 structures directly influence the formation of SEs.

Taken together, we propose a new model in which G4 structures regulate enhancer activity by influencing nucleosome occupancy and TF binding (Figure 7). The high enrichment of TFs, especially for architectural proteins including CTCF, RAD21, and SMC3, further facilitate the chromatin looping between G4 enhancers and their target promoters. The weak enhancers lack for G4 structures and displayed relatively high nucleosome occupancy, leading to low regulatory activity.

**Figure 7.**
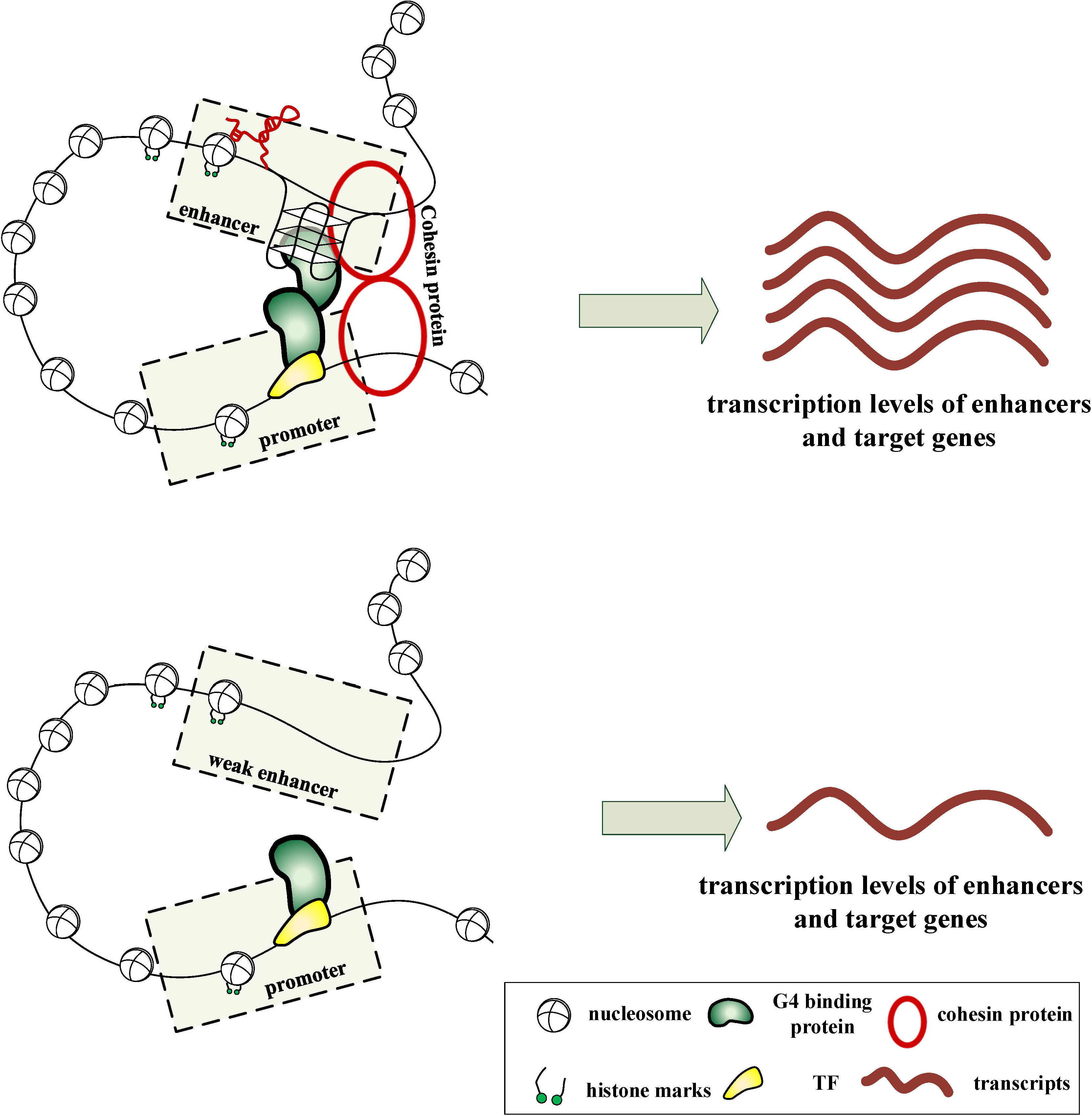
A schematic representation elucidating how G4 structures regulate the activity of enhancers. In the top panel, the formation of G4 structures on enhancers create an open chromatin environment for TFs to bind on, which lead to the transcription of enhancers. Abundant architectural proteins on G4 enhancers further facilitate enhancer-promoter interactions resulting in the high expression levels of target genes. In the bottom panel, there are no G4 structures on weak enhancers and the chromatin loop is unstable leading to relatively low expression levels of target genes.

Finally, we hypothesize that G4 structures may be a potential markers of enhancer regulatory activity. It can be seen that most G4 structures are rich in GRO-seq signals which has been proved to be indicators of active regulatory elements. Furthermore, we found that iG4s showed high regulatory activity even though they are not located in the known regulatory elements (Figure 6).

## Methods

### Identification of genomic elements

Human genes were downloaded from the GENCODE database (Harrow et al., 2012). The regions around TSSs (–3,000 bp to 3,000 bp) were designated as promoters. In the case of genes with more than one TSS, we selected the TSS closest to the 5’ end.

In line with previous research, H3K4me1 ChIP-seq peaks overlapped with DNase-seq peaks were designated as enhancers (Doni Jayavelu et al., 2020). The H3K4me1 ChIP-seq data of A549, HEK293T, HeLa-S3, and K562 cell lines were downloaded from GEO repository under accession number GSE91306, GSM3444906, GSM798322, and GSM788085, respectively. DNase-seq data of A549, HEK293T, HeLa-S3, and K562 cell lines were downloaded from ENCODE database under accession number ENCFF475BVB, ENCFF621ZJY, ENCFF775GBM, and ENCFF910QHN, respectively.

The downloaded public SRA files were processed by *fastq-dump* software. The fastq files were mapped to the human reference genome (GRCh37/hg19) by bowtie2 (Langmead and Salzberg, 2012). To identify ChIP-seq peak regions, we performed peak calling by MACS v2.0 with default parameters (Zhang et al., 2008). H3K4me1 peaks overlapped with DNase-seq peaks were used to define the enhancer regions, followed by further filtering based on the criteria: excluding H3K4me1 peaks overlapped with any previous defined promoters. SEs were identified by using the ROSE algorithm (Whyte et al., 2013) based on the H3K4me1 ChIP-seq signals with the default parameters.

### G4P and BG4 ChIP-seq data

G4P and BG4 ChIP-seq data were achieved from the GEO repository under accession number GSE133379 and GSE107690, respectively. The clean sequence reads were mapped to the human reference genome (GRCh37/hg19) by bowtie2 (Langmead and Salzberg, 2012). To identify exact G4 peaks, we performed peak calling by MACS v2.0 with default parameters (Zhang et al., 2008).

### ChIA-pet data

The original ChIA-PET data on the four cell lines (A549, HEK293T, HeLa-S3, and K562) were retrieved from the ENCODE website.

### GRO-seq data and RNA-seq data

The global nuclear run-on sequencing (GRO-seq) data of the four cell lines were generated by Bouvy-Liivrand et al. (GEO accession number: GSE92375) (Bouvy-Liivrand et al., 2017). GRO-seq are capable of capturing 5′-capped RNAs from active transcriptional regulatory elements with high accuracy (Danko et al., 2015). GRO-seq reads were mapped to the human reference genome (GRCh37/hg19) by Bowtie2 (Langmead and Salzberg, 2012).

The paired-end RNA-seq data were generated by Thomas Gingeras group of the ENCODE Consortium (Consortium, 2012). RNA-seq reads were mapped to the human reference genome (GRCh37/hg19) by tophat (Trapnell et al., 2012). We used cufflinks to generate transcriptome assembly (Trapnell et al., 2010).

### TF ChIP-seq Enrichment analysis

The genomic fold enrichment of TF binding profiles over G4 enhancers was determined using the ChIP-ATLAS enrichment analysis web tool (https://chip-atlas.org/enrichment_analysis) with the following parameters: antigen class: TFs and others; threshold for significance: 500. We selected the TFs with fold enrichment > 5 and q-value <1e-10 for each ‘antigen name’ as enriched TFs.

## Funding

This work was supported by the Shaanxi Province Science Foundation for Youth under grant number 2021JQ-023; the China Postdoctoral Science Foundation under grant number 2021M692581.

## Conflict of interest

The authors declare that they have no competing interests.

## Notes

### Competing Interest Statement

The authors have declared no competing interest.

## Reference

Agarwal, T., Roy, S., Kumar, S., Chakraborty, T.K., and Maiti, S. (2014). In the sense of transcription regulation by G-quadruplexes: asymmetric effects in sense and antisense strands. Biochemistry 53, 3711–3718.

Andersson, R., and Sandelin, A. (2020). Determinants of enhancer and promoter activities of regulatory elements. Nature reviews Genetics 21, 71–87.

Bouvy-Liivrand, M., Hernandez de Sande, A., Polonen, P., Mehtonen, J., Vuorenmaa, T., Niskanen, H., Sinkkonen, L., Kaikkonen, M.U., and Heinaniemi, M. (2017). Analysis of primary microRNA loci from nascent transcriptomes reveals regulatory domains governed by chromatin architecture. Nucleic acids research 45, 12054.

Cascon, A., and Robledo, M. (2012). MAX and MYC: a heritable breakup. Cancer research 72, 3119–3124.

Chambers, V.S., Marsico, G., Boutell, J.M., Di Antonio, M., Smith, G.P., and Balasubramanian, S. (2015). High-throughput sequencing of DNA G-quadruplex structures in the human genome. Nat Biotechnol 33, 877–881.

Chen, H., and Liang, H. (2020). A High-Resolution Map of Human Enhancer RNA Loci Characterizes Super-enhancer Activities in Cancer. Cancer cell 38, 701–715 e705.

Consortium, E.P. (2012). An integrated encyclopedia of DNA elements in the human genome. Nature 489, 57–74.

Core, L.J., Martins, A.L., Danko, C.G., Waters, C.T., Siepel, A., and Lis, J.T. (2014). Analysis of nascent RNA identifies a unified architecture of initiation regions at mammalian promoters and enhancers. Nature genetics 46, 1311–1320.

Danko, C.G., Hyland, S.L., Core, L.J., Martins, A.L., Waters, C.T., Lee, H.W., Cheung, V.G., Kraus, W.L., Lis, J.T., and Siepel, A. (2015). Identification of active transcriptional regulatory elements from GRO-seq data. Nature methods 12, 433–438.

De Santa, F., Barozzi, I., Mietton, F., Ghisletti, S., Polletti, S., Tusi, B.K., Muller, H., Ragoussis, J., Wei, C.L., and Natoli, G. (2010). A large fraction of extragenic RNA pol II transcription sites overlap enhancers. PLoS biology 8, e1000384.

Doni Jayavelu, N., Jajodia, A., Mishra, A., and Hawkins, R.D. (2020). Candidate silencer elements for the human and mouse genomes. Nature communications 11, 1061.

Du, Z., Zhao, Y., and Li, N. (2009). Genome-wide colonization of gene regulatory elements by G4 DNA motifs. Nucleic acids research 37, 6784–6798.

Ganji, M., Shaltiel, I.A., Bisht, S., Kim, E., Kalichava, A., Haering, C.H., and Dekker, C. (2018). Real-time imaging of DNA loop extrusion by condensin. Science 360, 102–105.

Georgakopoulos-Soares, I., Morganella, S., Jain, N., Hemberg, M., and Nik-Zainal, S. (2018). Noncanonical secondary structures arising from non-B DNA motifs are determinants of mutagenesis. Genome research 28, 1264–1271.

Hah, N., Murakami, S., Nagari, A., Danko, C.G., and Kraus, W.L. (2013). Enhancer transcripts mark active estrogen receptor binding sites. Genome research 23, 1210–1223.

Hansel-Hertsch, R., Beraldi, D., Lensing, S.V., Marsico, G., Zyner, K., Parry, A., Di Antonio, M., Pike, J., Kimura, H., Narita, M., et al. (2016). G-quadruplex structures mark human regulatory chromatin. Nature genetics 48, 1267–1272.

Hansel-Hertsch, R., Simeone, A., Shea, A., Hui, W.W.I., Zyner, K.G., Marsico, G., Rueda, O.M., Bruna, A., Martin, A., Zhang, X., et al. (2020). Landscape of G-quadruplex DNA structural regions in breast cancer. Nature genetics.

Hansel-Hertsch, R., Spiegel, J., Marsico, G., Tannahill, D., and Balasubramanian, S. (2018). Genome-wide mapping of endogenous G-quadruplex DNA structures by chromatin immunoprecipitation and high-throughput sequencing. Nature protocols 13, 551–564.

Harrow, J., Frankish, A., Gonzalez, J.M., Tapanari, E., Diekhans, M., Kokocinski, F., Aken, B.L., Barrell, D., Zadissa, A., Searle, S., et al. (2012). GENCODE: the reference human genome annotation for The ENCODE Project. Genome research 22, 1760–1774.

Henderson, E., Hardin, C.C., Walk, S.K., Tinoco, I., Jr., and Blackburn, E.H. (1987). Telomeric DNA oligonucleotides form novel intramolecular structures containing guanine-guanine base pairs. Cell 51, 899–908.

Holder, I.T., and Hartig, J.S. (2014). A matter of location: influence of G-quadruplexes on Escherichia coli gene expression. Chemistry & biology 21, 1511–1521.

Hong, S., and Kim, D. (2017). Computational characterization of chromatin domain boundary-associated genomic elements. Nucleic acids research 45, 10403–10414.

Huppert, J.L., and Balasubramanian, S. (2007). G-quadruplexes in promoters throughout the human genome. Nucleic acids research 35, 406–413.

Kim, T.K., Hemberg, M., Gray, J.M., Costa, A.M., Bear, D.M., Wu, J., Harmin, D.A., Laptewicz, M., Barbara-Haley, K., Kuersten, S., et al. (2010). Widespread transcription at neuronal activity-regulated enhancers. Nature 465, 182–187.

Lam, M.T., Li, W., Rosenfeld, M.G., and Glass, C.K. (2014). Enhancer RNAs and regulated transcriptional programs. Trends Biochem Sci 39, 170–182.

Langmead, B., and Salzberg, S.L. (2012). Fast gapped-read alignment with Bowtie 2. Nature methods 9, 357–359.

Li, L., Lyu, X., Hou, C., Takenaka, N., Nguyen, H.Q., Ong, C.T., Cubenas-Potts, C., Hu, M., Lei, E.P., Bosco, G., et al. (2015). Widespread rearrangement of 3D chromatin organization underlies polycomb-mediated stress-induced silencing. Mol Cell 58, 216–231.

Li, L., Williams, P., Ren, W., Wang, M.Y., Gao, Z., Miao, W., Huang, M., Song, J., and Wang, Y. (2020). YY1 interacts with guanine quadruplexes to regulate DNA looping and gene expression. Nature chemical biology.

Li, P.T., Wang, Z.F., Chu, I.T., Kuan, Y.M., Li, M.H., Huang, M.C., Chiang, P.C., Chang, T.C., and Chen, C.T. (2017). Expression of the human telomerase reverse transcriptase gene is modulated by quadruplex formation in its first exon due to DNA methylation. The Journal of biological chemistry 292, 20859–20870.

Li, W., Notani, D., Ma, Q., Tanasa, B., Nunez, E., Chen, A.Y., Merkurjev, D., Zhang, J., Ohgi, K., Song, X., et al. (2013). Functional roles of enhancer RNAs for oestrogen-dependent transcriptional activation. Nature 498, 516–520.

Li, W., Notani, D., and Rosenfeld, M.G. (2016). Enhancers as non-coding RNA transcription units: recent insights and future perspectives. Nature reviews Genetics 17, 207–223.

Mao, S.Q., Ghanbarian, A.T., Spiegel, J., Martinez Cuesta, S., Beraldi, D., Di Antonio, M., Marsico, G., Hansel-Hertsch, R., Tannahill, D., and Balasubramanian, S. (2018). DNA G-quadruplex structures mold the DNA methylome. Nature structural & molecular biology 25, 951–957.

Melgar, M.F., Collins, F.S., and Sethupathy, P. (2011). Discovery of active enhancers through bidirectional expression of short transcripts. Genome biology 12, R113.

Rao, S.S.P., Huang, S.C., Glenn St Hilaire, B., Engreitz, J.M., Perez, E.M., Kieffer-Kwon, K.R., Sanborn, A.L., Johnstone, S.E., Bascom, G.D., Bochkov, I.D., et al. (2017). Cohesin Loss Eliminates All Loop Domains. Cell 171, 305–320 e324.

Renciuk, D., Rynes, J., Kejnovska, I., Foldynova-Trantirkova, S., Andang, M., Trantirek, L., and Vorlickova, M. (2017). G-quadruplex formation in the Oct4 promoter positively regulates Oct4 expression. Biochim Biophys Acta 1860, 175–183.

Rodon, L., Svensson, R.U., Wiater, E., Chun, M.G.H., Tsai, W.W., Eichner, L.J., Shaw, R.J., and Montminy, M. (2019). The CREB coactivator CRTC2 promotes oncogenesis in LKB1-mutant non-small cell lung cancer. Science advances 5, eaaw6455.

Sanborn, A.L., Rao, S.S., Huang, S.C., Durand, N.C., Huntley, M.H., Jewett, A.I., Bochkov, I.D., Chinnappan, D., Cutkosky, A., Li, J., et al. (2015). Chromatin extrusion explains key features of loop and domain formation in wild-type and engineered genomes. Proceedings of the National Academy of Sciences of the United States of America 112, E6456–6465.

Sartorelli, V., and Lauberth, S.M. (2020). Enhancer RNAs are an important regulatory layer of the epigenome. Nature structural & molecular biology 27, 521–528.

Shen, J., Varshney, D., Simeone, A., Zhang, X., Adhikari, S., Tannahill, D., and Balasubramanian, S. (2021). Promoter G-quadruplex folding precedes transcription and is controlled by chromatin. Genome biology 22, 143.

Tang, Z., Luo, O.J., Li, X., Zheng, M., Zhu, J.J., Szalaj, P., Trzaskoma, P., Magalska, A., Wlodarczyk, J., Ruszczycki, B., et al. (2015). CTCF-Mediated Human 3D Genome Architecture Reveals Chromatin Topology for Transcription. Cell 163, 1611–1627.

Trapnell, C., Roberts, A., Goff, L., Pertea, G., Kim, D., Kelley, D.R., Pimentel, H., Salzberg, S.L., Rinn, J.L., and Pachter, L. (2012). Differential gene and transcript expression analysis of RNA-seq experiments with TopHat and Cufflinks. Nature protocols 7, 562–578.

Trapnell, C., Williams, B.A., Pertea, G., Mortazavi, A., Kwan, G., van Baren, M.J., Salzberg, S.L., Wold, B.J., and Pachter, L. (2010). Transcript assembly and quantification by RNA-Seq reveals unannotated transcripts and isoform switching during cell differentiation. Nature biotechnology 28, 511–515.

Varshney, D., Spiegel, J., Zyner, K., Tannahill, D., and Balasubramanian, S. (2020). The regulation and functions of DNA and RNA G-quadruplexes. Nature reviews Molecular cell biology.

Vietri Rudan, M., Barrington, C., Henderson, S., Ernst, C., Odom, D.T., Tanay, A., and Hadjur, S. (2015). Comparative Hi-C reveals that CTCF underlies evolution of chromosomal domain architecture. Cell Rep 10, 1297–1309.

Wang, X., Brandao, H.B., Le, T.B., Laub, M.T., and Rudner, D.Z. (2017). Bacillus subtilis SMC complexes juxtapose chromosome arms as they travel from origin to terminus. Science 355, 524–527.

Whyte, W.A., Orlando, D.A., Hnisz, D., Abraham, B.J., Lin, C.Y., Kagey, M.H., Rahl, P.B., Lee, T.I., and Young, R.A. (2013). Master transcription factors and mediator establish super-enhancers at key cell identity genes. Cell 153, 307–319.

Yang, Y., Su, Z., Song, X., Liang, B., Zeng, F., Chang, X., and Huang, D. (2016). Enhancer RNA-driven looping enhances the transcription of the long noncoding RNA DHRS4-AS1, a controller of the DHRS4 gene cluster. Sci Rep 6, 20961.

Zaug, A.J., Podell, E.R., and Cech, T.R. (2005). Human POT1 disrupts telomeric G-quadruplexes allowing telomerase extension in vitro. Proc Natl Acad Sci U S A 102, 10864–10869.

Zhang, Y., Liu, T., Meyer, C.A., Eeckhoute, J., Johnson, D.S., Bernstein, B.E., Nusbaum, C., Myers, R.M., Brown, M., Li, W., et al. (2008). Model-based analysis of ChIP-Seq (MACS). Genome biology 9, R137.

Zheng, K.W., Zhang, J.Y., He, Y.D., Gong, J.Y., Wen, C.J., Chen, J.N., Hao, Y.H., Zhao, Y., and Tan, Z. (2020). Detection of genomic G-quadruplexes in living cells using a small artificial protein. Nucleic acids research 48, 11706–11720.

